# Rapid and Active Stabilization of Visual Cortical Firing Rates Across Light-Dark Transitions

**DOI:** 10.1101/542670

**Authors:** Alejandro Torrado Pacheco, Elizabeth I. Tilden, Sophie M. Grutzner, Brian J. Lane, Yue Wu, Keith B. Hengen, Julijana Gjorgjieva, Gina G. Turrigiano

## Abstract

The dynamics of neuronal firing during natural vision are poorly understood. Surprisingly, mean firing rates of neurons in primary visual cortex (V1) of freely behaving rodents are similar during prolonged periods of light and darkness, but it is unknown whether this reflects a slow adaptation to changes in natural visual input, or insensitivity to rapid changes in visual drive. Here we use chronic electrophysiology in freely behaving rats of either sex to follow individual V1 neurons across many dark-light (D-L) and light-dark (L-D) transitions. We show that, even on rapid timescales (1s to 10 min), neuronal activity was only weakly modulated by transitions that coincided with the expected 12h/12h light-dark cycle. In contrast, a larger subset of V1 neurons consistently responded to unexpected L-D and D-L transitions, and disruption of the regular L-D cycle with 60 hours of complete darkness induced a robust increase in V1 firing upon re-introduction of visual input. Thus, V1 neurons fire at similar rates in the presence or absence of natural stimuli, and significant changes in activity arise only transiently in response to unexpected changes in the visual environment. Further, although mean rates were similar in L and D, pairwise correlations were significantly stronger during natural vision, suggesting that information about natural scenes in V1 is more readily extractable from correlations than from individual firing rates. Together, our findings show that V1 firing rates are rapidly and actively stabilized during expected changes in visual input, and are remarkably stable at both short and long timescales.

**Significance Statement:** The firing dynamics of neurons in primary visual cortex (V1) are poorly understood. Indeed, V1 neurons of freely behaving rats fire at the same mean rate in light and darkness. It is unclear how this stability is maintained, and whether it is important for sensory processing. We find that transitions between light and darkness happening at expected times have only modest effects on V1 activity. In contrast, both unexpected transitions and light re-exposure after extended darkness robustly increase V1 firing. Finally, pairwise correlations in neuronal spiking are significantly higher during the light, when natural vision is occurring. These data show that V1 firing is remarkably stable, and that neuronal correlations may represent sensory information better than mean firing rates.

## Introduction

Neurons in the cerebral cortex are spontaneously active, but the function of this internally generated activity is largely unexplained. Ongoing activity has been proposed to be noise due to random fluctuations (Zohary et al., 1994; Shadlen and Newsome, 1998; Averbeck et al., 2006). However other experiments have shown that spontaneous activity possesses coherent spatio-temporal structure (Arieli et al., 1995; Tsodyks et al., 1999; Ch’ng and Reid, 2010), suggesting it may play an important role in the processing of natural sensory stimuli (Arieli et al., 1995; Kenet et al., 2003; Fiser et al., 2004; MacLean et al., 2005; Luczak et al., 2009, 2013). In primary visual cortex (V1), spontaneous activity observed in complete darkness is similar to that evoked by visual stimulation with random noise stimuli, and is only subtly modulated by natural scene viewing (Gallant et al., 1998; Fiser et al., 2004). Recently we showed that individual V1 neurons have very stable mean firing rates in freely behaving rodents, and that these mean rates are indistinguishable in light and dark when averaged across many hours (Hengen et al., 2016). How V1 firing can be stable across such drastic changes in the visual environment while still meaningfully encoding sensory stimuli, and whether this stability is actively maintained or simply arises from intrinsic circuit dynamics, remains unknown.

Regulation of individual firing rates around a stable set point is thought to be essential for proper functioning of cortical circuits in the face of developmental or experience-dependent perturbations to connectivity (Miller and MacKay, 1994; Turrigiano and Nelson, 2004). Long-term stability of individual mean firing rates has now been observed in rodent V1 (Hengen et al., 2013, 2016; Keck et al., 2013) and M1 (Dhawale et al., 2017), suggesting it is a general feature of neocortical networks; further, perturbing firing rates in V1 through prolonged sensory deprivation results in a slow but precise homeostatic regulation of firing back to an individual set-point, showing that neurons actively maintain these set points over long time-scales (Hengen et al., 2016). This stability in mean firing rates, even across periods of light and dark, raises the question of how natural visual input is encoded by V1 activity in freely behaving animals. One possibility is that changes in visual drive result in rapid fluctuations in mean firing rates that operate over seconds to minutes. Another possibility is that firing rates are stabilized even over these short time scales, and visual information is primarily encoded in higher order network dynamics.

To generate insight into these questions we followed firing of individual neurons in V1 of freely behaving young rats over several days, as animals experienced normal light-dark (L-D) and dark-light (D-L) transitions, or transitions that were unexpectedly imposed. We found that expected L-D transitions had a very modest effect on firing rates of both excitatory and inhibitory neurons, even when examined immediately around the time of the transition. Population activity did not change significantly across these transitions, and when examined at the level of individual neurons only a small subset (~15%) of excitatory neurons consistently responded, and then only during D-L transitions when animals were awake. Interestingly, randomly timed transitions throughout the light-dark cycle elicited more consistent responses across sleep-wake states and at both D-L and L-D transitions, and robust and widespread responses to D-L transitions could be unmasked by exposing animals to prolonged darkness for 60 hours. These results suggest that the stability normally observed at expected (circadian) L-D transitions reflects an active process of stabilization. Finally, although mean rates were very similar in L and D, the pairwise correlations between simultaneously recorded neurons were significantly higher in the light than in the dark, even when controlling for behavioral state. Together our findings show that firing rates in V1 are actively stabilized as animals navigate dramatic changes in the visual environment, and that the correlational structure of V1 activity may carry more information about natural visual scenes than firing rates.

## Materials and Methods

All surgical and experimental procedures were approved by the Animal Care and Use Committee of Brandeis University and complied with the guidelines of the National Institutes of Health.

### *Surgery and* in vivo *experiments*

The data analyzed in this study were collected in previous electrophysiological recordings (Hengen et al., 2016; n = 7 rats), as well as from newly implanted animals (n = 7 rats). All surgical procedures were as described previously (Hengen et al., 2013). Briefly, Long-Evans rats of either sex were bilaterally implanted with custom 16-channel 33 μm tungsten microelectrode arrays (Tucker-Davis Technologies, Alachua, FL) into monocular primary visual cortex (V1m) on postnatal day 21 (P21). Location was confirmed post-hoc via histological reconstruction. Two EMG wires were implanted deep in the nuchal muscle. Animals were allowed to recover for two full days post-surgery in transparent plastic cages with *ad libitum* access to food and water. Recording began on the third day after surgery. The recording chamber (a 12”x12” plexiglass cage with walls lined with high contrast low spatial frequency gratings) was lined with 1.5” of bedding and housed two rats. Animals had *ad libitum* food and water, and were separated by a clear plastic divider with 1” holes to allow for tactile and olfactory interaction while preventing jostling of headcaps and arrays. Electrodes were connected to commutators (TDT) to allow animals to freely behave throughout the recordings. Novel toys were introduced every 24h, to promote activity and exploration. Lighting and temperature were kept constant (LD 12:12, lights on at 7:30 am, 21° C, humidity 25%-55%). Data were collected continuously for nine to eleven days (200-240 hours). Some animals (n = 11 rats) underwent a lid suture and/or eye reopening procedure on the third day of recording; in the present study we only analyzed data collected from the control hemisphere ipsilateral to the manipulated eye. For dark-exposure experiments, animals were kept in the dark starting on days 4 and 5 of the recording (i.e. starting at the time of lights off on day 3, from P26 to P28). Lights came on at the regular time (7:30 AM) on day 6 (P29).

### Electrophysiological recordings

*In vivo* electrophysiological recordings were performed as previously described (Hengen et al., 2016). Briefly, data were acquired at 25 kHz, digitized and streamed to disk for offline processing using a Tucker-Davis Technologies Neurophysiology Workstation and Data Streamer. Spike extraction and sorting was performed using custom MATLAB code. Spikes were detected as threshold crossings (−4 s.d. from mean signal) and re-sampled at 3x the original rate. Each wire’s waveforms were then subjected to principal component analysis (PCA) and the first four principal components were used for clustering using KlustaKwik (Harris et al., 2000). Clusters were merged or trimmed as described previously (Hengen et al., 2016). Spike sorting was done using custom MATLAB code relying on a random forest classifier trained on a manually scored dataset of 1200 clusters. For each cluster identified from the output of KlustaKwik, we extracted a set of 19 features, including ISI contamination (% of ISIs < 3 ms), similarity to regular spiking unit (RSU) and fast-spiking (FS) waveform templates, 60 Hz noise contamination, rise and decay time and slope of the mean waveform, waveform amplitude and width. Cluster quality was also ensured by thresholding of L-Ratio and Mahalanobis distance (Schmitzer-Torbert et al., 2005). The random forest algorithm classified clusters as noise, multi-unit or single-unit. Only single units with a clear refractory period were used for further analysis. Units were classified as RSU or FS based on the time between the negative peak and the first subsequent positive peak of the mean waveform (Fig. 1B). Clusters were classified as RSUs if this value was > 0.39 msec, and as FS otherwise (Niell & Stryker, 2008), with a lower threshold of 0.19 msec to eliminate noise. We used previously established criteria and methods to select neurons that we could reliably follow over time (Hengen et al., 2016); only neurons that could be recorded for at least 48 consecutive hours were used for analysis of light-dark transitions. For extended dark experiments, we analyzed all neurons that were online for at least one hour preceding and one hour following the time of light re-exposure. For estimates of mean firing during the extended dark phase, we analyzed the activity of all cells that could be recorded in the first and last 12-hour period during the 60 hours of darkness.

**Figure 1.**
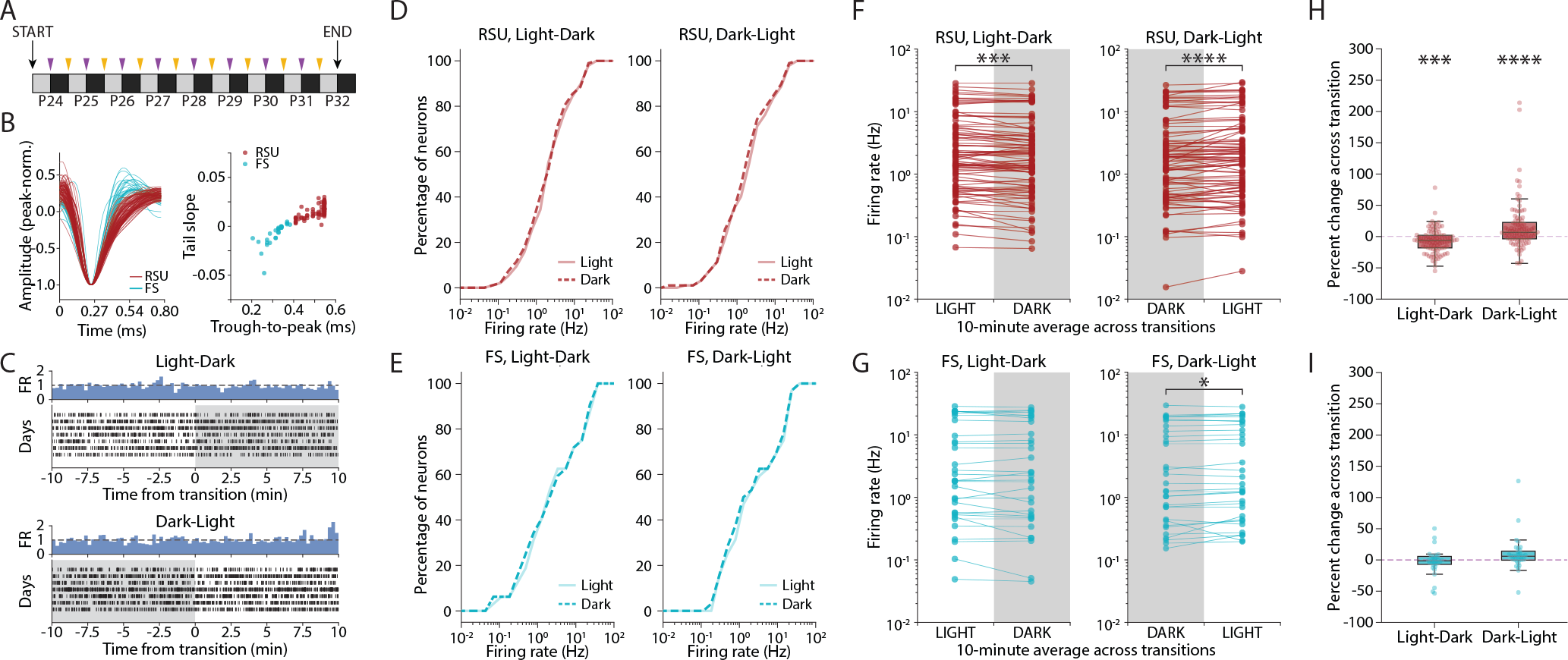
Circadian L-D and D-L transitions have a small effect on V1 firing rates. ***A***, Experimental protocol. Single-unit recordings were obtained from juvenile rats for a continuous 9-day period (P24-P32). Throughout this period animals were kept in a regular 12h/12h light/dark cycle and thus underwent light-dark (L-D, purple arrows) and dark-light (D-L, yellow arrows) transitions at regular 12-hour intervals. ***B***, Left, average waveform for each continuously recorded unit, identified as regular spiking unit (RSU, red) or fast-spiking cell (FS, blue). Right, plot of trough-to-peak time vs waveform slope 0.4 ms after trough reveals the bi-modal distribution used to classify recorded units as RSU or FS. ***C***, Example raster plot of spiking activity for a recorded unit across several days, showing 20 minutes of activity centered on the L-D (top) and D-L (bottom) transitions. Blue bars represent the peri-event time histogram obtained by averaging across days. ***D***, Cumulative distributions of RSU firing rates averaged over the 10 minutes of light (solid line) or dark (dashed line) around the transitions, for L-D (left) and D-L (right) transitions (L-D, p = 0.875; D-L, p = 0.99, two-sample Kolmogorov-Smirnov test). ***E***, As in D, but for FS units (L-D, p = 0.99; D-L p =1.0, two-sample Kolmogorov-Smirnov test). ***F***, Mean firing rate for each RSU, averaged across all transitions experienced by that neuron, in L-D (left) and D-L (right) transitions. Paired data indicates the average FR is for the same neuron. Distributions were not significantly different (L-D, p = 0.677; D-L, p = 0.655; Wilcoxon rank-sum test), but individual neurons across the whole distribution showed consistent changes at the transitions (L-D, *** p = 0.0002; D-L, **** p < 0.0001; Wilcoxon signed-rank test). ***G***, as in F, but for FS units. Distributions were not different (L-D, p = 0.905; D-L, p = 0.827; Wilcoxon rank-sum test), but individual FS units changed their firing consistently at D-L, but not L-D transitions (L-D, p = 0.318; D-L, * p = 0.026; Wilcoxon signed-rank test). ***H***, Percent change in firing rate across transition for RSUs (L-D, −7.09% ± 1.99%, *** p = 0.0006; D-L, 15.60% ± 4.00%, p = **** 0.0002; one-sample t-test). ***I***, as H, for FS units. Percentage change in FR was not different from 0 in either condition (L-D, −2.75% ± 3.80%, p = 0.475; D-L: 9.73% ± 5.12%, p = 0.067, one-sample t-test).

### Behavioral state classification

The behavioral state of animals was classified using a combination of local field potential (LFP), EMG and estimate of locomotion based on video analysis (see Hengen et al., 2016). LFPs were extracted from three separate recorded channels, resampled at 200 Hz, and averaged. The power spectral density was computed in 10-second bins using a fast Fourier transform method (MATLAB “spectrogram” function) using frequency steps of 0.1 Hz from 0.3 to 15 Hz. Power in the delta (0.3 - 4 Hz) and theta (6 – 9 Hz) bands was computed as a fraction of total power in each time bin. A custom algorithm was used to score each 10-second bin and assign one of four behavioral codes, based on the power in each frequency band as well as EMG and movement activity: active wake (high EMG and movement, low delta and theta), quiet wake (low EMG and movement, low delta and theta), REM sleep (very low EMG, no movement, low delta, high theta), and NREM sleep (low EMG and movement, high delta, low theta). For each animal, each hour of data was scored separately. The first ten hours were scored manually, and used as an initial training set for a random forest classifier (implemented in Python). The classifier was then used to score each successive hour, with manual corrections performed as needed. The classifier was re-trained after every hour scored, with a maximum number of 10,000 bins used for training (only the latest 10,000 bins were used).

### Extended darkness, immunostaining and image analysis

For analysis of c-fos protein following extended darkness, we transferred animals to custom dark box on P21. A light timer was set up to allow for complete control of the light-dark cycle inside the box. When animals were P26, lights were allowed to turn off at the regular time (7:30 PM), and set up to not turn back on. Animals were in complete darkness for 60 hours, from the night of P26 until night of P28 (ages matched with electrophysiological recordings). On the morning of P29, lights were allowed to turn back on at 7:30 AM. Animals were allowed to experience one hour of light before being deeply anesthetized and transcardially perfused. Control animals were either not exposed to darkness but kept on a regular 12h cycle, or anesthetized at 7:30 AM on P29 (before lights on) in the dark using night-vision goggles and then immediately perfused. Brains were fixed in 3.7% formaldehyde and 60 μm coronal slices of V1m were taken on a vibratome (Leica VT1000S). Slices were immersed in a solution of PBS and NaN_3_ and stored for immunostaining. To ensure consistent results between groups, all conditions were run in parallel. Slices were incubated for in a primary antibody solution (1:1000, Rabbit anti-c-fos, Cell Signaling Technologies) at room temperature for 24 hours. They were then rinsed and incubated for 2 hours with a secondary antibody (anti-rabbit Alexa Fluor 568, 1:400, Thermofisher). Sections were mounted on microscope slides with a DAPI-containing medium (DAPI Fluoromount-G, Southern Biotech), coverslipped and allowed to dry for 24 hours before imaging. Imaging was performed on a confocal microscope (Zeiss Laser Scanning Microscope 880). A 10X objective was used to take z-stacks of V1m in the DAPI and c-fos channels. Imaging settings were optimized for each staining/imaging session and kept constant across conditions; all conditions were imaged on a given session. Images were imported into Metamorph software for analysis. A granularity analysis was used to determine locations of cell bodies, and co-localized DAPI- and c-fos-positive granules were counted as c-fos-positive neurons. For each slice we analyzed the whole field of view, excluding the slice edges as they displayed DAPI staining artifacts.

### Analysis of electrophysiological data

All electrophysiology data was analyzed using a custom code package written in Python. The precise time of lights on/off was determined by analysis of video recordings or using a light-sensitive resistor. All analyses were performed on the 10 minutes before and after transitions. Peri-event time histograms were obtained by binning data in 0.25 second bins and normalizing data to the pre-transition period. Firing rates were estimated by sliding a 1 or 2 minute window in 20-second steps. Mean and s.e.m. were estimated by averaging across days. To compare population data across transitions we calculated the average firing rate in the 10 minutes before and after the transition without binning. For analysis restricted to a given behavioral state, we only considered transitions during which the animal was in that state for the whole 20 minutes (10 minutes before and after the transition). To estimate the number of individual neurons that consistently changed their firing rate in response to L-D and D-L transitions, we used a paired t-test to determine whether the neuron’s firing followed a consistent pattern of change across multiple transitions. We used a bootstrap method to estimate the number of cells expected to pass our significance threshold by chance; for each iteration of the bootstrap, we chose a random time point within the first 24h. We then created dummy transition times at 12 hour intervals from that starting time point, and used these dummy transition points to repeat the above analysis for each cell. This procedure was repeated 100 times (i.e. with 100 different dummy transition points), to obtain 100 values for the percent of significantly changing cells. We used this dataset to estimate the mean and 95% confidence intervals for this parameter. Only neurons that were followed through at least 4 transitions were used for analysis of circadian transitions. For non-circadian transitions, we analyzed neurons that experienced at least 6 transitions.

### Pairwise correlations

Each spike train was binned into spike counts of bin size 100 ms, generating a vector of spike counts for each cell. The spike count correlation coefficient ρ for a pair of neurons was computed in 30 minute episodes using a sliding window of 5 minutes. This produced 139 values for each neuron pair on every single half day (12 hours of light and 12 hours of darkness). The average of these values then determined the correlation value of each pair for every single half day:

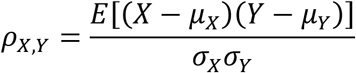

where *X* and *Y* represent the spike count vectors of two cells, respectively; *μ*_*X*_ and *μ*_*Y*_ are the means of *X* and *Y*; σ_*X*_ and σ_*Y*_ denote the standard deviations of *X* and *Y*; *E* is the expectation. This produced the matrices of pairwise spike count correlations on different half days. To generate the normalized correlation curve, correlations were normalized to the average correlation of each animal at P26 during the light period. Correlations in mixed behavioral states were computed with the above-stated method using the entire 12-hour periods of L or D, while correlations in wake only took into account the wake episodes. Results with bin size of 5 ms followed the same approach.

### Experimental design and statistical analysis

For paired data, both for firing rates and spike count correlations, comparisons were done using a Wilcoxon signed-rank test. To compare a population mean to a given value (e.g. 0) we used one-sample t-tests. To compare cumulative distributions we used Kolmogorov-Smirnov tests. Data are represented as mean ± s.e.m. Box plots represent median ± interquartile range, with whiskers extending to the rest of the distribution. All statistical analyses were performed using Python.

### Code accessibility

All code used for analysis is available from the authors upon request.

## Results

Neurons in V1 maintain remarkably similar mean firing rates in L and D, but how L-D transitions affect firing on more rapid time scales in freely viewing and behaving animals is unclear. Here we use chronic *in vivo* electrophysiological recordings from freely behaving rats to closely examine the activity of V1 neurons at D-L and L-D transitions in regular (12h/12h) and manipulated light/dark cycles, and during unexpected light-dark transitions. Using previously established methods (Hengen et al., 2016) we followed individual neurons over time and across multiple light transitions. This approach allowed us to analyze the dynamics of neuronal activity at different timescales in response to the appearance or disappearance of natural visual stimuli.

### The appearance or disappearance of natural visual stimuli has only a modest effect on the mean firing rates of V1 neurons

The firing rates of V1 neurons recorded in freely behaving young rats in light and dark are exceedingly similar when averaged in 12-hour periods (Hengen et al., 2016). Here we combine previously and newly acquired datasets and set out to re-analyze the activity of V1 neurons around the transition from presence to absence of visual input (light-dark, L-D), and vice versa (dark-light, D-L) (Fig. 1A). Recorded neurons were classified as regular spiking units (RSU, n = 96) or fast-spiking cells (FS, n = 32) based on waveform shape and according to established criteria (Fig. 1B; Niell & Stryker, 2008; Hengen et al., 2013). These populations are mostly comprised of excitatory pyramidal neurons (RSU) and GABAergic parvalbumin-containing interneurons (FS) (Kawaguchi & Kubota, 1993; Nowak et al., 2003).

As rats experience L-D or D-L transitions, most neurons showed little change in firing (Fig. 1C). We treated each transition as a separate trial and estimated the firing rate for each cell as the average of the peri-event time histogram (PETH) centered on the transition. We first aimed to compare activity at the population level in different stimulus conditions. To this end, we determined whether the distributions of mean firing rates averaged over 10 minutes on either side of the L-D and D-L transitions were similar to each other. Cumulative distributions in light and dark were indistinguishable, for both RSU and FS cells, in all conditions (Fig. 1D, E; two-sample Kolmogorov-Smirnov test, RSU, L-D: p = 0.88; D-L: p = 0.99; FS, L-D: p = 0.99; D-L: p = 1.0). Similarly, when we compared the distributions using a Wilcoxon rank-sum test, we found no difference between the distributions of mean firing rates before vs. after the transitions (Wilcoxon rank-sum test, RSU, L-D: p = 0.677; D-L: p = 0.655; FS, L-D: p = 0.905; D-L: p = 0.827).

Next we took advantage of our ability to follow individual neurons across transitions to examine the data in a paired manner, where the firing rate of each neuron was compared before and after the transition. For each neuron we computed mean firing rate in the 10 minutes before and after the transition time, and averaged across transitions of the same type to estimate the average effect on individual neuronal firing. This analysis revealed a small but consistent change in mean RSU firing rates across both L-D and D-L transitions (Fig. 1F; Wilcoxon sign-rank test: L-D: p = 0.0002; D-L: p < 0.0001), while the activity of FS cells only changed significantly at D-L transitions (Fig. 1G; Wilcoxon sign-rank test: L-D: p = 0.318; D-L: p = 0.026). The magnitude of these effects was small, on order 7-15% for RSUs (Fig. 1H, I; RSU, L-D: −7.09% ± 1.99%, p = 0.0006; D-L: 15.60% ± 4.00%, p = 0.0002; FS, L-D: −2.75% ± 3.80%, p = 0.475; D-L: 9.73% ± 5.12%, p = 0.067, one-sample t-test).

These data show that, surprisingly, dramatic changes in visual input cause very minor changes in V1 firing rates. The distributions of mean rates in the presence and absence of natural visual stimuli are identical in the proximity of transitions. Analysis of many transitions shows that RSU firing rates are consistently affected when the visual environment changes, but this modulation is decidedly modest.

### Behavioral state affects sensitivity of firing rates to visual stimuli

As rats were freely behaving throughout all experiments, we considered whether their alertness state at the light transitions could affect the activity of V1 neurons. LFP, EMG and video data were collected and used to score animals’ behavioral state into either asleep or awake (Hengen et al., 2016). For each animal, 20-minute periods centered on the L-D and D-L transitions were considered. Only periods during which the animal remained in the same behavioral state for the entire time were analyzed. For each neuron, we plotted the mean firing rate before the transition against the mean rate after the transition. The activity of neurons proved to be strikingly similar across all transitions, whether the animals were awake or asleep (Fig. 2A, B). In either behavioral state, firing rates in light and dark were very strongly correlated, and the slope of the regression line was close to one (RSU, wake, L-D: slope = 0.959, *r* = 0.966, p < 10^−43^; D-L: slope = 0.960, *r* = 0.991, p < 10^−48^; FS, wake, L-D: slope = 1.113, *r* = 0.964, p < 10^−13^; D-L: slope = 0.976, *r* = 0.990, p < 10^−18^; RSU, sleep, L-D: slope = 1.147, *r* = 0.941, p < 10^−12^; D-L: slope = 0.990, *r* = 0.996, p < 10^−13^; FS, sleep, L-D: slope = 1.093, *r* = 0.987, p < 10^−13^; D-L: slope = 1.003, *r* = 0.996, p < 10^−11^).

**Figure 2.**
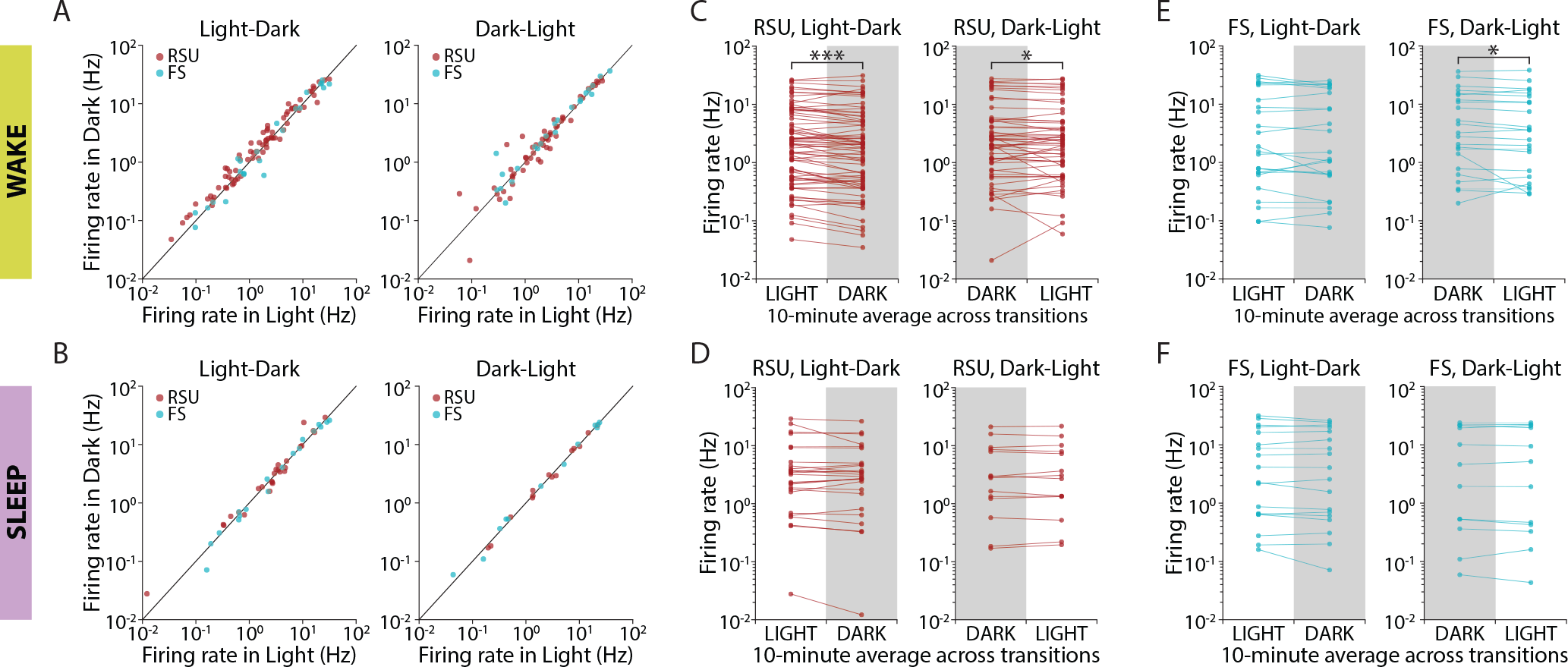
Changes in natural visual input modestly modulate the firing of V1 neurons during wake but not sleep. ***A***, Comparison of mean firing rates, in 10-minute averages around the transitions, in L-D and D-L transitions when the animal was awake for the whole 20 minutes. Activity in light and dark was strongly correlated for both transition types. ***B***, as in A, but for transitions during which animals were asleep for the 20 minute period around the transition. Firing rates in light and dark during sleep were also strongly correlated. ***C***, Mean firing rate of individual RSUs, calculated in 10-minute averages around luminance transitions and averaged across all transitions during which animals were awake. Neuronal activity changed consistently at the transitions (L-D, p = 0.0001; D-L, p = 0.0457; Wilcoxon signed-rank test). ***D***, as in C, but for transitions during which animals were asleep. No significant change was observed (L-D, p = 0.656; D-L, p = 0.925; Wilcoxon signed-rank test). ***E***, As in C, for FS units. Cells’ activity only changed significantly at D-L transitions (L-D, p = 0.689; D-L, p = 0.039; Wilcoxon signed-rank test). ***F***, As in D, for FS cells. No significant change was observed (L-D, p = 0.557; D-L, p = 0.638; Wilcoxon signed-rank test).

We again looked at the data in paired form, by comparing a neuron’s average firing rate on either side of a L-D transition. The mean activity of RSUs in V1 changed consistently across transitions when animals were awake (Fig. 2C; L-D: p = 0.0001; D-L: p = 0.0457, Wilcoxon signed-rank test), but not when they were asleep (Fig. 2D; L-D: p = 0.656; D-L: p = 0.925, Wilcoxon signed-rank test). We observed a similar pattern in FS cells, although the data in the wake condition was not significant for light-dark transitions (Fig. 2E, F; Wake, L-D: p = 0.689; D-L: p = 0.039; Sleep, L-D: 0.557; D-L: p = 0.638; Wilcoxon signed-rank test). Once again, these effects were of small magnitude (7-12%). Thus, V1 neurons do not respond to expected (circadian) changes in the visual environment when animals are asleep, and respond only modestly when animals are awake.

### A sub-population of RSUs is consistently responsive to dark-light transitions

While we only detected small changes at the population levels (and no change in the population distribution), we occasionally observed neurons whose activity appeared to be consistently modulated by visual stimuli. The majority of neurons showed no spiking modulation across multiple transitions (Fig. 3A), but a subset of neurons showed higher activity on one side of the transition (Fig. 3B). Occasionally neurons responded to both L-D and D-L transitions (Fig. 3B), but more often neurons were only responsive to one or the other. To quantify these observations, we treated each transition independently for each neuron, and averaged firing rates for 10 minutes before and after lights on/off, and identified neurons that changed their firing rate consistently across transitions using a paired t-test.

**Figure 3.**
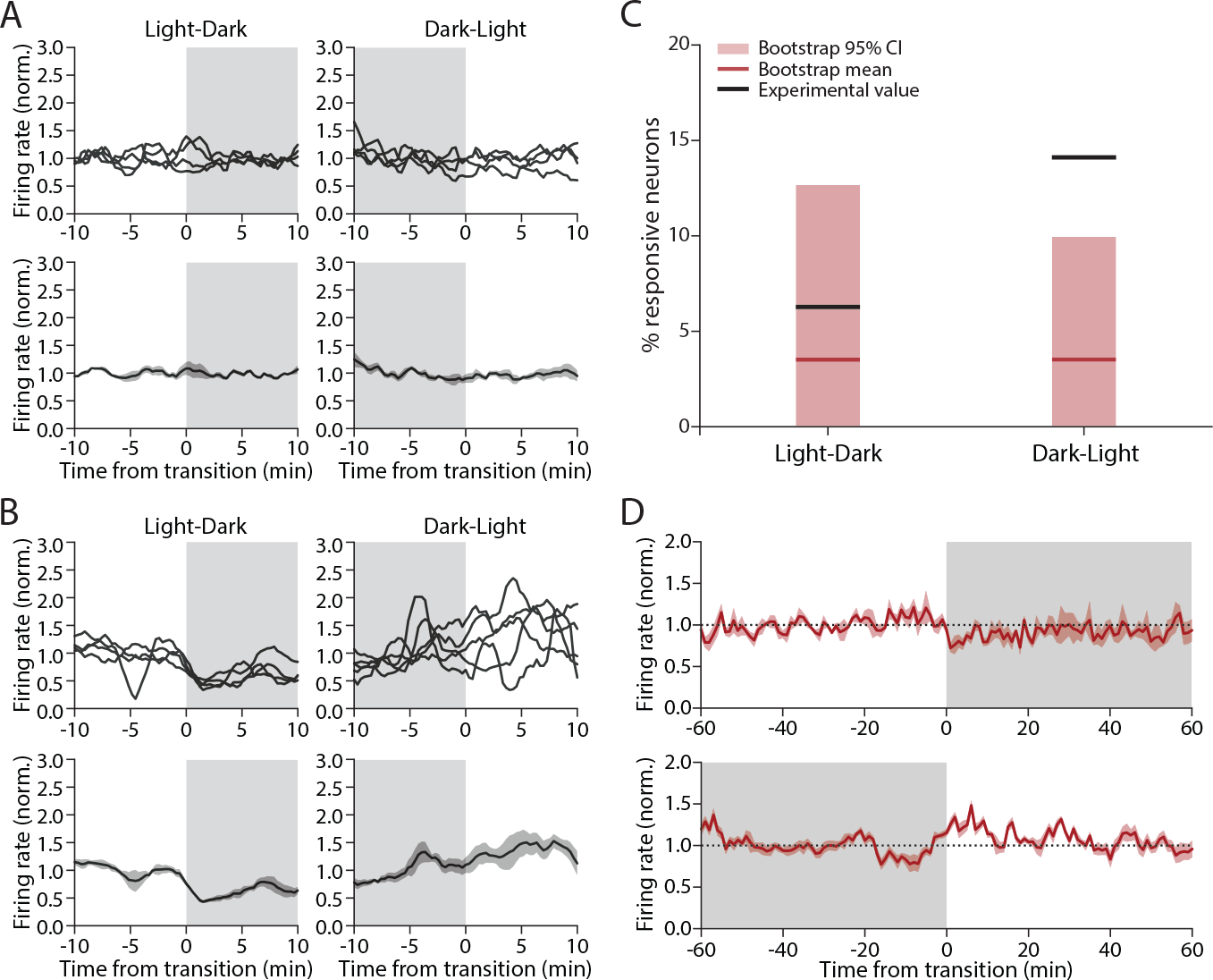
A subset of RSUs consistently increase their firing rate in response to expected dark-light transitions. ***A***, example of a RSU unresponsive to light transitions (left, L-D; right, D-L). Top, binned firing rate for each transition; bottom, average across transitions. ***B***, example RSU that responds consistently to luminance transitions. ***C***, Percentage of RSUs found to be responsive to L-D (left) and D-L (right) transitions and bootstrap control. Black lines show experimental value (actual % of responsive neurons); red line shows bootstrap mean; light red bar shows extent of the bootstrap 95% confidence interval (L-D, actual value: 6.25%, bootstrap mean: 3.55%; 95% CI: 0% - 12.62%; D-L, actual value: 14.06%; bootstrap mean: 3.09%; 95% CI: 0% - 9.91%; n = 64). ***D***, Mean firing rate averaged across transitions for all D-L-responsive RSUs, calculated for two hour around each transition for L-D (top) and D-L (bottom) transitions. The transient nature of the firing rate response is visible in the bottom panel.

Because neuronal firing rates are variable, we presumed that some of these apparent responses were spurious. To estimate the false positive rate we performed a bootstrap analysis using random time points as dummy “transitions”. We chose nine transition points 24 hours apart from each other (to match circadian transitions) and analyzed mean firing rates for each neuron as above but using these dummy transition points. This process was repeated 100 times to arrive at an estimate of the mean and 95% confidence interval for the percentage of responsive cells (mean and 95% CI, RSU, L-D: 3.55% [0% - 12.62%], D-L: 3.09% [0% - 9.91%]; n= 64).

The proportion of cells we found to be transition-responsive was within the range expected by chance for all conditions except for RSUs in D-L transitions (Fig. 3C). We found that 14% of RSUs in our experimental condition had significantly changing firing rates from dark to light, well outside the range expected by chance (95% CI for this group: [0% - 9.91%]). In addition, most of these neurons (88.9%) showed an increase in firing rate at the onset of light, while in the bootstrap control neurons were found to have an equal probability of increasing or decreasing their activity at a given transition point (51.8% of neurons increasing).

Finally, we examined the temporal dynamics of firing rate changes for the subset of RSUs that were consistently responsive to D-L transitions. We plotted the mean activity within 1 hour of the transition, across all transitions and across neurons for this subpopulation (Fig. 3D). On average, the change in FR was short-lived, on the order of ~10 minutes, and of moderate size (~25% increase). This analysis shows that a small subset of excitatory pyramidal neurons in V1 consistently modulate their activity in response to the expected appearance of visual input after a circadian 12-hour period of darkness. This change is transient, with firing rates returning to pre-transition levels within minutes.

### Light transitions have no effect on average ISIs over short timescales

Our analysis so far shows that, on a timescale of 10s of minutes, few V1 neurons show significant firing rate modulation to the appearance or disappearance of natural visual stimuli. One possible explanation for this apparent lack of responsiveness is that these dramatic sensory changes trigger a rapid adaptation mechanism that quickly restores average V1 activity back to baseline. Such adaptation mechanisms within V1 have been well described, and can operate on a time-scale of 100s of ms to many minutes (Kohn, 2007; Wissig and Kohn, 2012; Benucci et al., 2013). To address this possibility, we examined neuronal firing in 1-, 10- and 30-second intervals around L-D and D-L transitions. Spiking in these short time windows was sparse and variable across days (Fig. 4A, B). We averaged the mean inter-spike interval (ISI) across days for each cell, and compared averages in the 10 seconds before and after transitions. To ensure we were not missing effects on even shorter timeframes we also computed the mean ISI in 1-second windows around the transitions. For both the 1-sec and the 10-sec case, we found no statistically significant effect (Fig. 4C, 1-sec, L-D: p = 0.27; D-L: p = 0.36; Fig. 4D, 10-sec, L-D: p = 0.97; D-L: p = 0.31; Wilcoxon sign-rank test). Similar results were obtained when this analysis was carried out with 5-sec and 30-sec intervals (data not shown). This indicates that the stability of firing across transitions is not due to a short-term adaptation process that rapidly restores firing to baseline.

**Figure 4.**
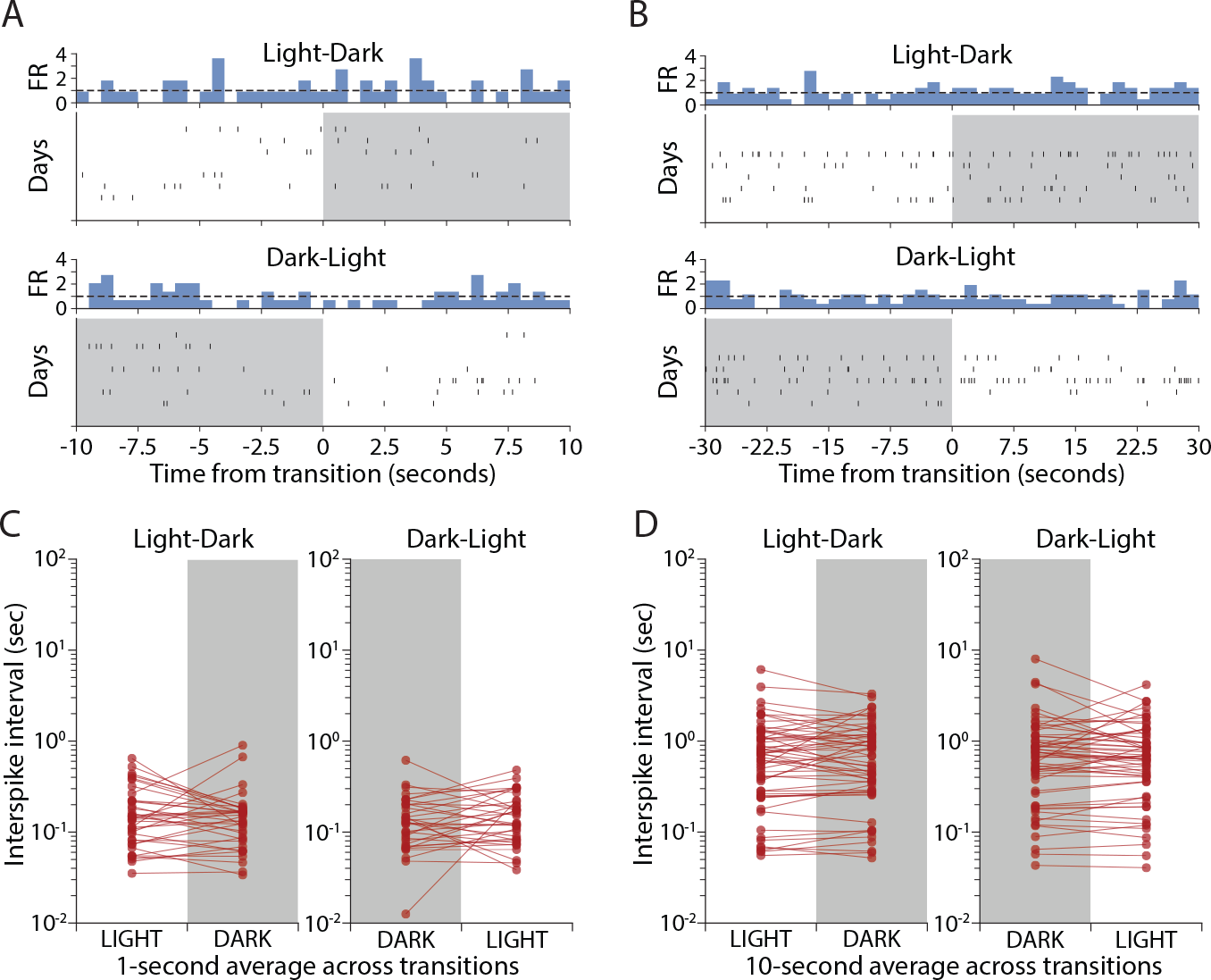
L-D and D-L transitions have no effect on V1 firing rates over short timescales. ***A***, Raster plot showing activity of one example RSU in a 10-second interval around L-D and D-L transitions. Vertical ticks represent spikes, rows represent transitions happening on different recording days. ***B***, A second example RSU, showing a 30-second interval around transitions. ***C,*** Mean ISI for all recorded cells, obtained by averaging ISIs in 10-second bins around L-D and D-L transitions for different days. Each dot represents the mean for one cell, obtained by averaging across days. No significant change was observed (L-D, p = 0.97; D-L, p = 0.31; Wilcoxon sign-rank test). ***D***, As in C, but for 1-second averages. No significant change was observed (L-D, p = 0.27; D-L, p = 0.36; Wilcoxon sign-rank test).

### Non-circadian, unexpected L-D transitions are more likely to perturb V1 firing rates

All of our data so far suggest that dramatic changes in visual input at circadian L-D transitions have very subtle effects on V1 firing. We wondered if this might be due to circadian entrainment, i.e. that when L-D and D-L transitions happen at regular times they are expected and the response of neurons to otherwise salient stimuli is attenuated. To test this, we examined neuronal responses to stimulus transitions occurring at random points in the circadian cycle.

We recorded single-unit activity in V1 while turning the lights off (or on) for 10 minutes during the light (or dark) cycle (Fig. 5A, n = 6 animals). We then calculated the number of neurons that consistently and significantly changed their firing at these unexpected transitions, and again used a bootstrap analysis to calculate the false positive rate. In marked contrast to expected transitions (Fig. 3D), we found that both L-D and D-L unexpected transitions caused a subset of RSUs to consistently modulate their spiking (Fig. 5B, C). This effect was seen regardless of behavioral state (significantly changing RSUs, sleep, L-D: 21.9%, n = 64; D-L: 13.4%, n = 67; wake, L-D: 17.6%, n = 91; D-L: 12.7%, n = 55) and the proportion of significantly changing neurons was higher than expected by chance in most conditions (bootstrap mean and 95% CI, RSU, sleep, L-D: 4.42% [0% - 8.95%], D-L: 4.31% [0% - 9.38%]; RSU, wake, L-D: 4.22% [0% - 8.79%], D-L: 4.33% [0% - 9.09%]).

**Figure 5.**
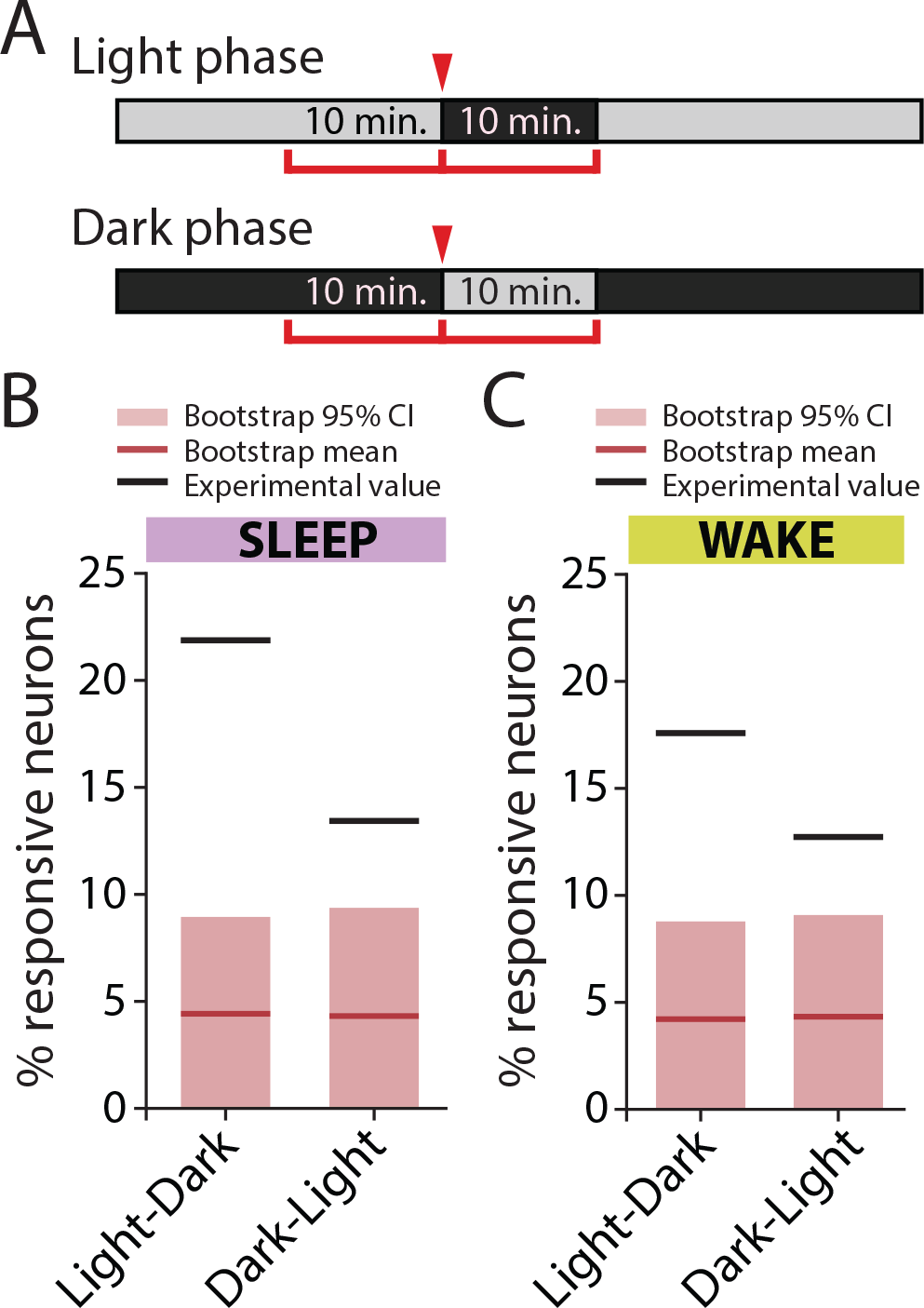
Randomly timed L-D and D-L transitions induce consistent firing rate changes in RSUs. ***A***, Experimental design. Animals were exposed to 10-minute periods of darkness during the light phase, and 10-minute periods of light during the dark phase, at random points throughout the light/dark cycle. Mean firing rates were calculated in 10-minute intervals around the transition. ***B,*** Percentage of RSUs that were responsive to luminance transitions, when transitions happened in epochs of sleep. Black line shows actual experimental value; red line shows bootstrap mean; light red bar covers the bootstrap 95% confidence interval (sleep, L-D: 21.9%, bootstrap mean and 95% CI: 4.42% [0% - 8.95%], n=64; D-L: 13.4%, bootstrap mean and 95% CI: 4.31% [0% - 9.38%], n=67). ***C***, as in B, but for transitions happening while animals were awake (wake, L-D: 17.6%, bootstrap mean and 95% CI: 4.22% [0% - 8.79%], n=91; D-L: 12.7%, bootstrap mean and 95% CI: 4.33% [0% - 9.09%], n=55).

These results show that more neurons respond consistently to L-D and D-L transitions when these do not line up with the circadian cycle the animals are entrained on. However, even during unexpected transitions only a minority (12%-20%) of neurons consistently changed their firing rate in response to the appearance or disappearance of natural visual stimuli.

### Pairwise correlations in V1 are significantly higher in light than in dark

To investigate whether higher order network properties are modified by the presence or absence of natural visual stimuli, we examined the structure of pairwise correlations in light and in dark (Fig. 6, n = 5 animals). Plotting the correlation matrices of one animal at P27 revealed that these correlations were higher in the light (calculated over the 12-hour period at P27) than in the dark (calculated over the 12-hour period at P27.5; Fig. 6A). We then plotted the average correlation computed continuously over 4 days (normalized to the average correlation of each animal at P26 in light) (Fig. 6B). The normalized pairwise correlation showed a pronounced oscillation across light and dark periods, and was consistently higher in the light. To assess the degree to which correlation of individual pairs changed, we compared the correlation of 922 pairs in light versus dark computed for spike counts with bin sizes of 5 or 100 ms, respectively. We found that correlations in light were higher than in dark for both bin sizes (Fig. 6C; left, 5 ms: p < 10^−70^; right, 100 ms: p < 10^−125^, Wilcoxon signed-rank test). To ensure that the observed difference of correlations between light and dark was not caused by disproportionate time spent in wake or sleep, we restricted the analysis to periods of wake, and again computed the average correlation. Consistent with our previous analysis, correlations in wake during light were significantly greater than in wake during dark (Fig. 6D; left, 5 ms: p < 10^−55^; right, 100 ms: p < 10^−110^, Wilcoxon signed-rank test). These results indicate that the presence of natural visual stimuli increases pairwise correlations in V1.

**Figure 6.**
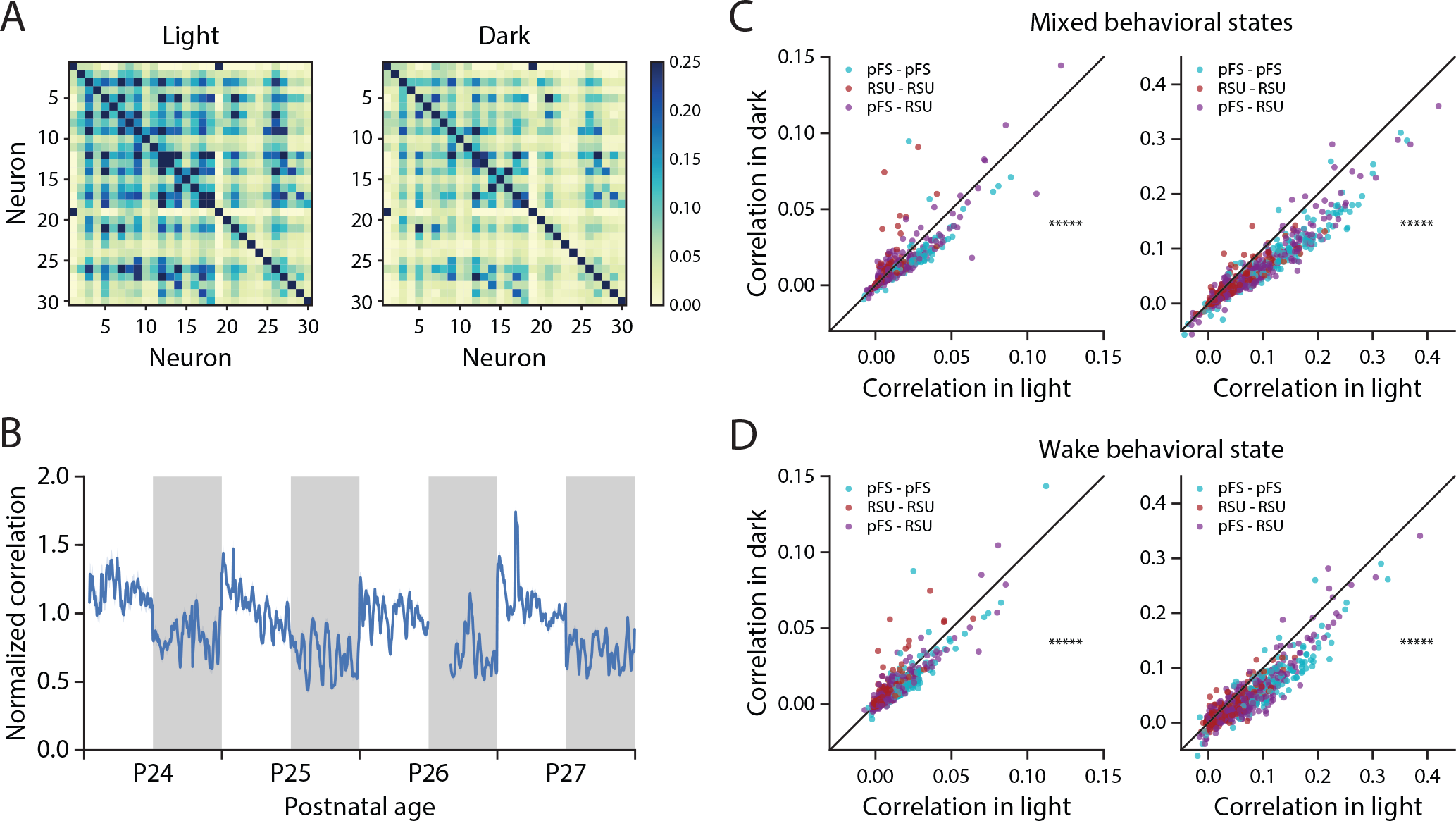
Pairwise correlations in V1 are higher during light than during dark. ***A***, Example pairwise correlation structure of 30 neurons from a single animal during light (left; calculated over the 12-hour period at P27) and during dark (right; calculated over the 12-hour period at P27.5). ***B***, The average correlation of 922 pairs from five animals over 4 days, normalized to the average correlation of each animal relative to P26 in light. The gap in the data at P26 corresponds to the time animals were anesthetized for monocular deprivation, which was excluded from analysis. ***C***, Comparison of the average correlation of 922 pairs during light and during dark with bin size 5ms (left) and bin size 100ms (right). (left: p < 10^-70; right: p < 10^-125, Wilcoxon signed-rank test). ***D***, Comparison of average correlation of 922 pairs in wake during light and during dark with bin size 5ms (left) and bin size 100ms (right). (left: p < 10^-55; right: p < 10^-110, Wilcoxon signed-rank test).

### Prolonged dark exposure enhances the responsiveness of V1 neurons to natural visual input

Our data show that re-exposure to light after 12 h of darkness has only modest effects on V1 firing; in contrast, re-exposing animals to light after a period of *prolonged* darkness is a standard paradigm for increasing activity-dependent gene expression in V1 (Rosen et al., 1992; Mower, 1994; Kaminska et al., 1996; reviewed in Kaczmarek and Chaudhuri, 1997). We therefore wondered whether prolonged dark exposure might unmask robust responses to the sudden onset of visual stimuli within V1.

We began by using expression of the immediate early gene *c-fos*, which is driven by enhanced calcium influx during elevated activity (Bartel et al., 1989; Sheng et al., 1990; reviewed in Sheng and Greenberg, 1990). After prolonged darkness, brief light exposure induces widespread *c-fos* expression in V1 of cats and rodents (Rosen et al., 1992; Kaplan et al., 1996; Yamada et al., 1999; Mower and Kaplan, 2002). To replicate this we placed P26 rats in the darkness for 60 hours (12 hours + 2 days) and then exposed them to light for one hour before immunostaining for the c-fos protein (light exposed, n = 28 slices, 5 animals). We used age-matched animals either exposed to one hour of light after a regular 12h/12h cycle (regular control, n = 22 slices, 4 animals), or kept in the dark for 60 hours but sacrificed before lights on (dark control, n = 23 slices, 4 animals), as controls (Fig. 7A, B). Animals in the light exposure condition showed an elevated percentage of c-fos-positive cells (Fig. 7C; RC: 11.4% ± 1.6%; DC: 6.1% ± 0.8%; LE: 16.8% ± 1.7%. LE vs RC p=0.032; LE vs DC p=0.001, one-way ANOVA with Tukey post-hoc test), as well as increased total staining intensity (Fig. 7D; normalized to RC, RC: 1.00 ± 0.06; DC: 0.79 ± 0.05; LE: 1.31 ± 0.09. LE vs RC p=0.011; LE vs DC p=0.001, one-way ANOVA with Tukey post-hoc test). These data confirm that a 60-hour period of prolonged darkness is sufficient to induce increased expression of *c-fos* in rodent V1.

**Figure 7.**
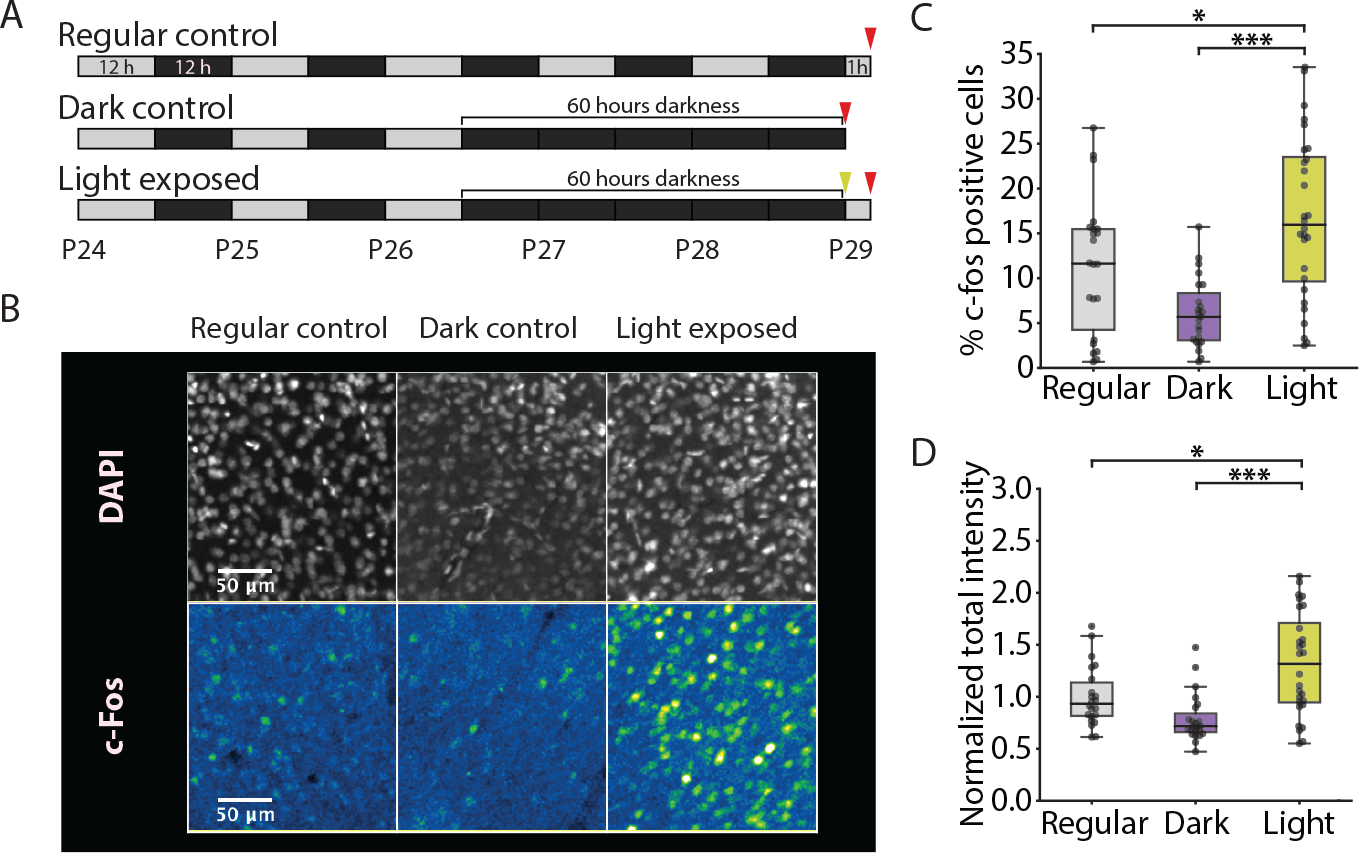
Prolonged darkness results in increased *c-fos* expression upon light re-exposure. ***A***, Experimental protocol. Animals were exposed to either a regular 12h/12h light/dark cycle, and sacrificed 1 hour after lights on at P29 (regular control, RC, n = 22 slices, 4 animals); exposed to darkness for 60 hours starting at P26 and sacrificed before lights on (dark control, DC, n = 23 slices, 4 animals); exposed to 60 hours of darkness starting at P26 and sacrificed 1 hour after light re-exposure (light exposed, LE, n = 28 slices, 5 animals). ***B***, representative images showing DAPI (top row) and *c-fos* (bottom row) immunostaining for regular control (left), dark control (middle), and light exposed (right) animals. ***C***, Percentage of *c-fos-*positive cells in all three groups (RC: 11.4% ± 1.6%; DC: 6.1% ± 0.8%; LE: 16.8% ± 1.7%. * p=0.032; *** p=0.001, one-way ANOVA with Tukey post-hoc test). ***D***, Total co-localized DAPI and c-fos staining intensity, normalized to average of RC group (RC: 1.00 ± 0.06; DC: 0.79 ± 0.05; LE: 1.31 ± 0.09; * p=0.011; *** p=0.001, one-way ANOVA with Tukey post-hoc test).

Next we asked whether this elevated *c-fos* expression was correlated with increased firing. We used the same paradigm as above but recorded continuously from V1 during the baseline, dark-exposure, and light re-exposure period (n = 4 animals). Upon light re-exposure, both RSUs and FS cells showed a substantial transient increase in firing rate at the time of lights on (Fig 8A; RSU: n = 32; FS: n = 12). We compared average firing rates 10 minutes before and after the transition for each cell. Both FS and RSU populations showed a significant increase in firing rate following light exposure (Fig. 8C, D; RSU: p < 10^−5^; FS: p = 0.034; Wilcoxon signed-rank test). The percent change in firing rate across the transition was also significantly different from zero (Fig. 8B; all cells: 87.1% ± 13.5%, p < 10^−^7; RSU: 80.7% ± 14.9%, p < 10^−5^; FS: 104.3% ± 29.8%, p = 0.005; one-sample t-test), and the majority of neurons increased their activity at lights on (RSU: 31/32 neurons; FS: 10/12 neurons).

**Figure 8.**
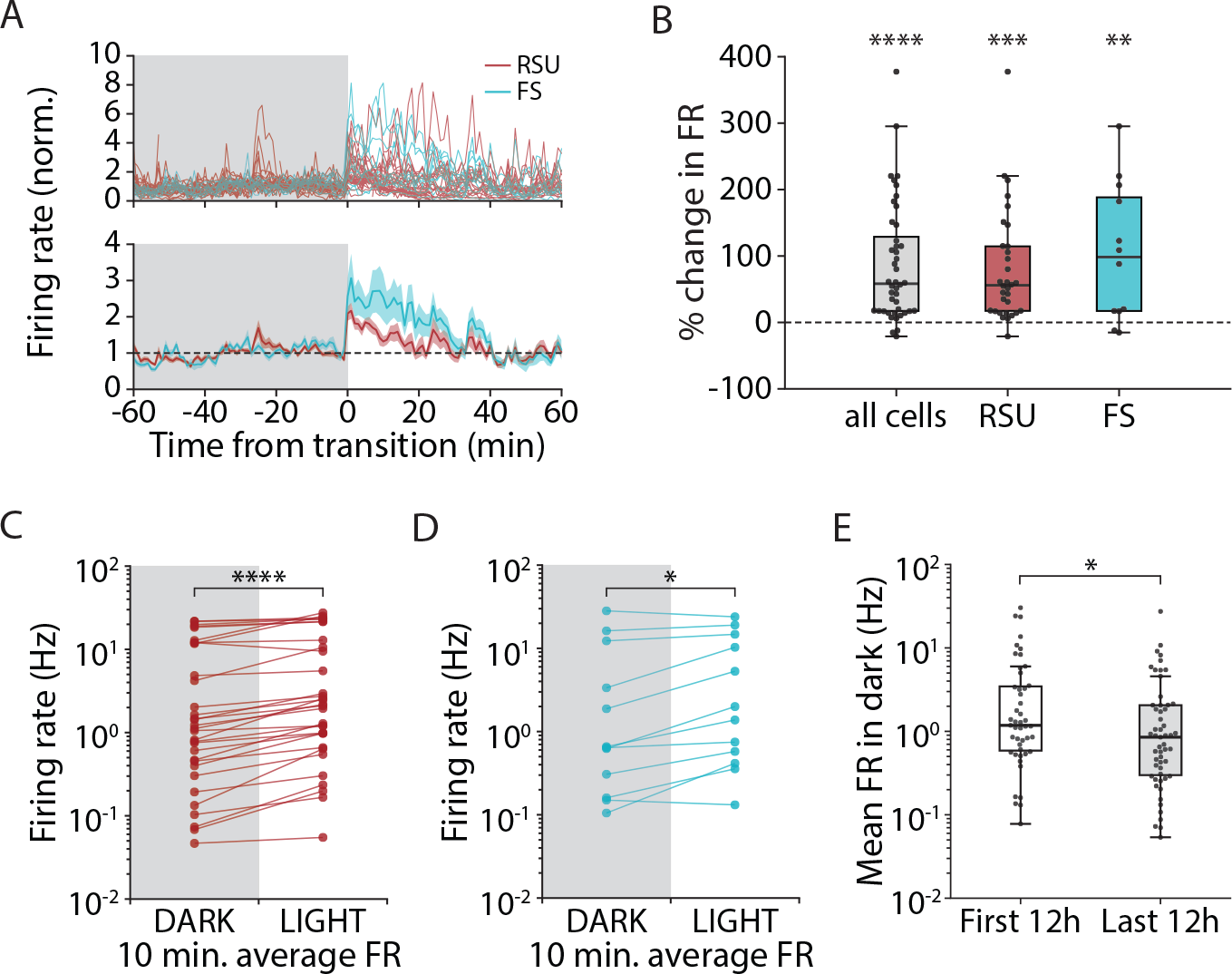
Light re-exposure after prolonged darkness causes a robust and widespread increase in V1 firing, following a slight reduction in firing rates over the dark period. ***A***, Time course of RSU and FS spiking in a 2-hour period around the time of light re-exposure. Individual unit traces (top) and average across cells (bottom) shows marked increase in firing at the time of lights on. ***B***, Percentage change in firing rate between the 10 minutes prior to and the 10 minutes immediately following light re-exposure (all cells, 87.1% ± 13.5%, n=44, **** p < 10^−7^; RSU, 80.7% ± 14.9%, n=32, *** p < 10^−5^; FS, 104.3% ± 29.8%, n=12, ** p = 0.005; one-sample t-test). ***C***, Mean firing rate in the 10 minutes before and after the transition, for each recorded RSU (**** p < 10^−5^, Wilcoxon signed-rank test). ***D***, As in C, for recorded FS cells (* p = 0.034, Wilcoxon signed-rank test). ***E***, Mean firing rate for all recorded RSUs, averaged across the first 12-hour period within the 60 hours of darkness (First 12h, mean ± s.e.m: 4.00 ± 0.97 Hz, median: 1.18 Hz, n = 47) and the last 12-hour period of darkness before light re-exposure (Last 12h, mean ± s.e.m: 2.27 ± 0.57 Hz, median: 0.85 Hz, n = 55; * p = 0.044, Wilcoxon rank-sum test).

It has been reported that prolonged dark exposure increases firing rates in rodent V1 (Bridi, de Pasquale, Lantz et al., 2018), suggesting that enhanced responsiveness to light re-exposure might arise from increased excitability of V1 circuitry. To examine this we asked how prolonged dark exposure affected RSU firing rates in freely behaving animals prior to light re-exposure. When we compared the distribution of RSU firing rates during the first and last 12 hours of the 60-hour long period of prolonged darkness, rather than an increase, we found a small but significant *decrease* in firing rates (Fig. 8E; mean ± s.e.m, first 12h: 4.00 ± 0.97 Hz, last 12h: 2.27 ± 0.57 Hz; median, first 12h: 1.18 Hz; last 12h: 0.85 Hz; p = 0.044, Wilcoxon rank-sum test). Thus the enhanced responsiveness to restoration of natural visual stimuli is unlikely to be due to a simple global increase in circuit excitability. These data indicate that prolonged dark exposure disrupts the normal stability of V1 firing across D-L transitions, and suggests that the maintenance of this stability is dependent upon visual experience.

## Discussion

How internal and external factors influence the long-term dynamics of neuronal firing in V1 is poorly understood. Here we recorded from ensembles of single units over a period of several days in freely viewing and behaving animals and found that firing rates of both excitatory and inhibitory V1 neurons were remarkably stable even when sensory input changed abruptly and dramatically. During expected circadian L-D transitions very few V1 neurons significantly changed their firing. A larger subset of V1 neurons were consistently responsive to unexpected L-D transitions, and disruption of the regular L-D cycle with two days of complete darkness induced a widespread and robust increase in V1 firing upon subsequent re-introduction of visual input. These data show that V1 neurons fire at similar rates in the presence or absence of natural visual stimuli, and that significant changes in mean activity arise only in response to unexpected changes in the visual environment. While mean firing rates were not different in L and D, pairwise correlations were significantly stronger in the light in the presence of natural visual stimuli, suggesting that information about natural scenes in V1 is more readily extractable from pairwise correlations than from individual firing rates. Taken together, our findings are consistent with a process of rapid and active stabilization of firing rates during expected changes in visual input, and demonstrate that firing rates in V1 are remarkably stable at both short and long timescales.

The near absence of firing rate modulation in response to the appearance (or disappearance) of natural visual stimuli may seem surprising, as there is a rich literature supporting the idea that V1 neurons respond to optimal stimuli by increasing their spiking (Hubel and Wiesel, 1959, 1962; Campbell et al., 1968; Pettigrew, 1974; Henry et al., 1974; Movshon, 1975; De Valois et al., 1982; Gizzi et al., 1990; Carandini and Ferster, 2000). Many of these studies used anesthetized preparations, making comparisons with our results difficult, but our data are consistent with previous reports of small differences in overall activity between natural vision and complete darkness in awake animals (Fiser et al., 2004), and sparse modulation of spiking in response to natural scene viewing (Gallant et al., 1998; Vinje and Gallant, 2000; Haider et al., 2010; Herikstad et al., 2011). In general our data support the view that mean firing rates in V1 can be stabilized over both long (Hengen et al., 2016) and short timescales without interfering with visual coding, which may arise through very sparse modulation of spiking and/or higher order population dynamics (Haider et al., 2010; Vinck et al., 2015; Dipoppa et al., 2018).

Despite the lack of robust changes in population spike rates at D-L transitions, we did observe a small subset of neurons that transiently increased their firing specifically at the appearance of visual input (Fig. 4). Since this occurred in freely behaving animals, it is unlikely that these neurons were responding to the same optimal visual stimuli on consecutive days. The more parsimonious explanation seems to be that these are broadly tuned neurons that become activated at most D-L transitions. Interestingly, we detected a greater proportion of such transition-responsive cells when light transitions happened randomly throughout the L-D cycle, including a population of neurons that transiently responded to non-circadian L-D transitions by decreasing their firing rate (Fig. 4). Thus unexpected changes in visual drive unmask robust and bidirectional changes in firing in a small subset (15-20%) of V1 neurons. There are several potential explanations for this effect. It is possible that the responsive neurons are specialized to represent this “unexpectedness” as an error signal, as has been proposed in some models of predictive coding (Rao and Ballard, 1999; Egner et al., 2010). Alternatively, it could be the result of modulation by other brain areas that encode the surprise signal, akin to that seen in response to attention or reward cues (Shuler and Bear, 2006; Stănişor et al., 2013), or during modulation of V1 by locomotion (Niell and Stryker, 2010). Finally, our data cannot distinguish between this last possibility and the opposite signal, i.e. a suppression of responses at expected transitions.

We were able to disrupt the normal conservation of firing rates across D-L transitions even more dramatically by using a prolonged dark-exposure paradigm, which induced a network-wide enhancement of firing upon light re-exposure. This paradigm is thought to induce metaplastic changes within V1 that increase AMPA quantal amplitude onto L2/3 pyramidal neurons (Goel and Lee, 2007; Blackman et al., 2012; Bridi, de Pasquale, Lantz et al., 2018), but the impact of these changes on overall V1 function and excitation/inhibition balance are unclear. A previous study in anesthetized animals found that several days of dark exposure increased firing rates in V1, raising the possibility that prolonged dark exposure increases overall V1 excitability (Bridi, de Pasquale, Lantz et al., 2018); however, here we found a small but significant reduction in mean firing rate across the population in freely behaving animals, suggesting that circuit excitability is if anything reduced by prolonged dark exposure. Although the circuit mechanism by which dark exposure unmasks robust responses to D-L transitions is unclear, these experiments suggest that normal visual experience is necessary to maintain the ability of V1 circuits to stabilize their firing across these transitions.

In contrast to our observations on the stability of firing rates, we found that pairwise correlations in visual cortex were markedly higher in the light phase than in the dark phase (Fig. 5). This is consistent with previous reports that ongoing spontaneous activity in the dark is less correlated than activity elicited by natural scene stimuli (Fiser et al., 2004; Karimipanah et al., 2017). Correlations are dependent on the degree of synchrony within neuronal circuits (Harris and Thiele, 2011; Schölvinck et al., 2015) and are higher during anesthesia (Greenberg et al., 2008), raising the possibility that this is a simple reflection of time spent in different behavioral states during the L and D phase. However, we observed the same increased correlation in L when only analyzing periods when animals were awake, ruling out this possibility. Thus, we conclude that, in freely behaving and viewing animals, sensory input can shift visual cortical circuits to more correlated dynamical states, even in a condition of low synchrony when animals are awake.

Our results add to a growing body of work suggesting that ongoing activity in mammalian V1 plays an important role in modulating sensory responses, as well as in integrating other sensory, motor, and motivational signals (Tsodyks et al., 1999; Rao and Ballard, 1999; Treue, 2001; Fiser et al., 2004; Luczak et al., 2009, 2013; Goard and Dan, 2009; Ringach, 2009; Egner et al., 2010; Niell and Stryker, 2010; Destexhe, 2011; Keller et al., 2012; Ayaz et al., 2013; Saleem et al., 2013; Vinck et al., 2015). Our results also show that firing rates of most V1 neurons are remarkably stable over both long and short time scales and in the presence and absence of visual information, suggesting that most visual information during natural viewing is not encoded by changes in firing rates. Instead, our data suggest that perturbations in firing primarily occur during unexpected changes in visual input, indicating an effect of entrainment/expectation and the existence of an active mechanism for stabilization of activity. This may be of particular importance given the observation that pairwise correlations are increased when animals are exposed to visual input, as global fluctuations in firing rate can strongly affect the strength of correlations between pairs of neurons (Harris and Thiele, 2011). Thus, it is possible that stable firing rates enable changes in correlations to reflect differences in sensory input, and hence to promote effective sensory processing.

## Acknowledgements

This work was supported by R01EY025613 (G.G.T. and A.T.P.) and the Max Planck Society (Y.W. and J.G.). The authors declare no competing financial interests. We thank Lauren Kronheim for experimental help.

